# FITM2 deficiency results in ER lipid accumulation, ER stress, reduced apolipoprotein B lipidation, and VLDL triglyceride secretion *in vitro* and in mouse liver

**DOI:** 10.1101/2023.12.05.570183

**Authors:** Haizhen Wang, Cyrus Nikain, Jaime Amengual, Maxwell La Forest, Yong Yu, Meng C. Wang, Russell Watts, Richard Lehner, Yunping Qiu, Min Cai, Irwin J. Kurland, Ira J. Goldberg, Sujith Rajan, M. Mahmood Hussain, Jeffrey L. Brodsky, Edward A. Fisher

## Abstract

**Objectives:** Triglyceride (TG) association with apolipoprotein B100 (apoB100) serves to form very low density lipoproteins (VLDL) in the liver. The repertoire of factors that facilitate this association is incompletely defined. FITM2, an integral endoplasmic reticulum (ER) protein, was originally discovered as a factor participating in cytoplasmic lipid droplets (LDs) in tissues that do not form VLDL. We hypothesized that in the liver, in addition to promoting cytosolic LD formation, FITM2 would also transfer TG from its site of synthesis in the ER membrane to nascent VLDL particles within the ER lumen.

**Methods:** Experiments were conducted using a rat hepatic cell line (McArdle-RH7777, or McA cells), an established model of mammalian lipoprotein metabolism, and mice. FITM2 expression was reduced using siRNA in cells and by liver specific cre-recombinase mediated deletion of the *Fitm2* gene in mice. Effects of FITM2 deficiency on VLDL assembly and secretion *in vitro* and *in vivo* were measured by multiple methods, including density gradient ultracentrifugation, chromatography, mass spectrometry, simulated Raman spectroscopy (SRS) microscopy, sub-cellular fractionation, immunoprecipitation, immunofluorescence, and electron microscopy.

**Main findings:** 1) FITM2-deficient hepatic cells *in vitro* and *in vivo* secrete TG-depleted VLDL particles, but the number of particles is unchanged compared to controls; 2) FITM2 deficiency in mice on a high fat diet (HFD) results in decreased plasma TG levels. The number of apoB100-containing lipoproteins remains similar, but shift from VLDL to LDL density; 3) Both *in vitro* and *in vivo*, when TG synthesis is stimulated and FITM2 is deficient, TG accumulates in the ER, and despite its availability this pool is unable to fully lipidate apoB100 particles; 4) FITM2 deficiency disrupts ER morphology and results in ER stress.

**Principal conclusions:** The results suggest that FITM2 contributes to VLDL lipidation, especially when newly synthesized hepatic TG is in abundance. In addition to its fundamental importance in VLDL assembly, the results also suggest that under dysmetabolic conditions, FITM2 may be a limiting factor that ultimately contributes to non-alcoholic fatty liver disease (NAFLD) and steatohepatitis (NASH).

## Introduction

Lipid droplets (LDs) are found in most eukaryotic cells and in some prokaryotes and span a wide range of sizes, from tens of nanometers to several microns in diameter. Nevertheless, the primary components of an LD universally consist of a core of neutral lipids—triglycerides (TGs), and sterol esters (SEs)—surrounded by a phospholipid monolayer[1]. LDs serve as reservoirs for lipids typically used for short-term energy generation and membrane synthesis; other functions include the release of fatty acids by adipocytes in response to fasting and protection from lipotoxicity in many tissues (reviewed in[1]). They also play important roles in cellular protein degradation[2, 3], the endoplasmic reticulum (ER) stress response[4, 5], and viral replication[6].

The ER is the major site of lipid synthesis in eukaryotic cells, and consequently LDs are generated in this organelle (reviewed in[7, 8]). Once synthesized, TG begins to accumulate within the bilayer of the ER membrane and forms a lens-like structure[1]. A simple model of LD formation is that once sufficient TG accumulates within the bilayer, the budding of a nascent LD into the cytosol occurs. The budding processes is catalyzed by myriad ER-associated factors whose identities and functions are still being characterized. Yet, once formed by the concerted action of these factors, mature LDs may completely separate from the ER, or—at least in yeast—some LDs can remain connected to the ER by a membrane bridge[9–12].

In contrast to other cell types (such as adipocytes), LD-resident TG in hepatocytes can provide both a source of short-term energy as well as serve as a lipid reservoir for the formation of nascent very-low density lipoproteins (VLDLs)[13]. These apolipoprotein B (apoB)-containing particles transport endogenous lipids to extrahepatic tissues. Although the factors that regulate the distribution of TG between LDs and VLDLs are not fully established, association of lipid with forming VLDL particles has been previously shown to involve MTTP[14],PLIN2[15], CIDEB[15], LPCAT3[16], apoC3[17], seipin[18] and Tm6sf2[19–21]. More recently, evidence has emerged to add the fat-storage-inducing transmembrane (FIT) proteins to this list (this BioRxiv report and [5]), which in higher eukaryotes are known as FITM1 and FITM2[10, 22].

The FITM proteins reside in the ER membrane and directly interact with TG to mediate LD formation in many cell types[9, 22]. Oxidative tissues, such as skeletal and cardiac muscle, express FITM1, whereas FITM2 is expressed ubiquitously, including adipose tissue and the liver[22]. The two homologs are ∼35% identical, and their overexpression favors the formation of LDs[23], albeit with somewhat different properties. One study reported that the absence of FITM2 in murine adipocytes leads to lipodystrophy[24]. Its overexpression in murine skeletal muscle or plants favors intracellular lipid accumulation and LD expansion[22, 25, 26]. Until recently, FITM2 was believed to regulate the budding and scission of LDs by altering monolayer tension in the ER[27]. However, FITM2 also possesses acyl CoA diphosphatase activity, and its absence alters ER morphology in cultured breast cancer cells[28]. In addition, FITM2 was proposed to directly regulate TG loading onto nascent VLDLs[29].

Here, we directly investigate this hypothesis by studying the role of hepatic FITM2 in the partitioning of newly-synthesized TG between the ER, LDs, and nascent VLDL particles *in vitro* and *in vivo*. We show that hepatic FITM2 deficiency results in increased intracellular lipid content, which is accompanied by reduced TG secretion in both cultured hepatocytes and liver-specific FITM2-deficient mice. Biochemical analyses reveal that FITM2 depletion reduces the lipidation of apoB100, the apolipoprotein onto which neutral lipids, sterols, sterol esters, and phospholipids assemble to form VLDLs[30]. Together, these data demonstrate that a FIT family member regulates hepatic lipid secretion.

## Methods

### Cell Culture

McArdle RH7777 rat hepatocytes (McA cells) stably expressing human apoB100[31] were grown in culture plates coated with type I collagen from calf skin (Sigma Aldrich, St. Louis, MO) as previously described[32] and cultured in growth media composed of Dulbecco’s modified – Eagle’s medium (DMEM, Corning, Corning, NY) containing 10% fetal bovine serum (FBS; Gemini, West Sacramento, CA), 10% horse serum (Gemini, West Sacramento, CA), 1% L-glutamine (Cellgro; Manassas, VA), and penicillin (100 U/ml)/streptomycin (100 ug/ml) (Sigma Aldrich). To stimulate LD formation and VLDL lipidation, lipid synthesis was enhanced with 0.6 mM oleic acid (OA) complexed to fatty acid-free BSA (0.12 mM) at a molar ratio of 5:1[33].

### siRNA Transfections

McA cells at 40% confluence were incubated with rat *Fitm2* siRNA (Dharmacon, Waltham MA), or untargeted control siRNA (Dharmacon) using DharmaFECT 4 Transfection Reagent in Opti-MEM media (Life Technologies, Carlsbad, CA), according to manufacturer’s protocol. Cells were maintained in normal growth media for 72 h, at which point they reached 80-90% confluence. FITM2 knock down efficiency was assessed by RT-PCR (see below for details).

### [^35^S]-Protein and [^14^C]-TG Labeling

McA cells were incubated with either OA complexed to fatty acid-free BSA as above, or BSA alone, in experimental medium (0.5% FBS, 0.5% horse serum, antibiotics). After a 2h incubation, cells were incubated in the presence of 120 μCi/ml of [^35^S]-protein labeling mix (Perkin-Elmer) or 0.25 μCi/ml of [^14^C]-glycerol (Perkin-Elmer) for 4 h. Conditioned media were collected and centrifuged at 500xg for 5 min, and cells were incubated with RIPA lysis buffer (50 mM Tris, pH 7.4, 150 mM NaCl, 0.25% sodium deoxycholate, 1% Nonidet P-40) in the presence of a protease inhibitor cocktail and phenylmethanesulfonyl fluoride (Sigma-Aldrich, St. Louis, MO).

### Immunoprecipitation and Quantification of [^35^S]-Labeled Proteins

ApoB100 quantification was performed as published (e.g.,[32]). Briefly, to immunoprecipitate radiolabeled apoB (apoB100 or apoB48), cell lysate or conditioned medium was mixed with 5 μg of goat anti-apoB100 (MilliporeSigma, Burlington, MA), NET buffer (150 mM NaCl, 5 mM EDTA, 50 mM Tris [pH7.4], 1% Triton X-100, and 0.1% SDS), and protein A Sepharose beads (Life Technologies). The mixture was incubated overnight at 4°C with gentle shaking. After incubation, the Sepharose beads were harvested, washed, and the immunoprecipitated proteins were released by the addition of SDS-PAGE sample buffer, followed by heating at 95°C for 5 min. Samples were resolved with 4% urea gels via SDS-PAGE, fixed in the presence of acetic acid and methanol, and dried. The dried gels were exposed to a phosphorimager plate, and the resulting images were used for the quantification of labeled proteins. Gel loading was normalized to total labeled protein obtained after precipitation of an aliquot with trichloroacetic acid (TCA).

### Density Gradient Ultracentrifugation

Cell media were loaded at the bottom of a centrifuge tube (Beckman Coulter Life Sciences, Indianapolis, IN) and overlaid by various volumes of KBr solutions at increasingly lower densities, as described previously[34]. The tubes were centrifuged at 34,600 rpm in a TH-641 swinging bucket rotor (ThermoFisher Scientific, Scientific, Waltham, MA) for 20h at 15 °C. After centrifugation, 10-11 fractions were collected, which contained distinct lipoproteins at the indicated densities, and apoB100 contents in the fractions were quantified by immunoprecipitation/SDS-PAGE as described just above.

### Triglyceride (TG) Mass Measurements

To measure TG content in McA cells, cell monolayers were washed with cold phosphate buffered saline (PBS), lysed with RIPA buffer (ThermoFisher), scraped, and then sonicated. The TG content was then quantified using a colorimetric method assay kit (Wako Diagnostics, Mountain View, CA). Plasma TG from mice were measured directly by the same kit. In plasma samples separated by FPLC, TG was similarly measured from each lipoprotein fraction (VLDL, LDL, HDL).

To measure TG content in mouse livers, ∼30 mg of frozen liver was homogenized in 300 µL RIPA buffer on ice. Lysates were sonicated and centrifuged at 8000xg for 1 min at room temperature to eliminate cell debris. Total TG content was then measured from the resulting supernatant as above.

### Lipid Isolation and Quantification of [^14^C]-TG

Radiolabeled TG was isolated and quantified as done previously[35]. Briefly, the lipid content of cell lysate or conditioned media was extracted using a mixture of Dole’s solvent (isopropanol: heptane: 0.5 M sulfuric acid; 40:10:1), heptane, and water. The extract was evaporated and dried using nitrogen gas and dissolved in chloroform. Lipid extracts were next resolved using thin layer chromatography (TLC) on a silica gel matrix with a mobile phase composed of hexane, diethyl ether, and acetic acid (80:20:2). Glyceryl trioleate (Sigma) was used as a TG standard. Lipid species were identified after staining with iodine vapors, and the individual species were scraped and mixed with scintillation fluid (Perkin-Elmer). [^14^C]-TG content was quantified using a LS6500 Beckman liquid scintillation counter (Beckman Coulter, LS6500).

### Animals and Husbandry

All animal procedures and experiments were approved by the New York University Institutional Animal Use and Care Committee. *Fitm2* floxed (*Fitm2^fl/fl^*, JAX stock #028832) and Albumin-Cre (*Alb-cre^+/+^*, JAX stock #003574) mice were purchased from the Jackson Laboratory (Bar Harbor, ME)[24, 36]. Liver-specific *Fitm2* knockout mice (*Fitm2^fl/fl^* / *Alb-cre^+/+^*) were generated by crossing *Fitm2^fl/fl^* and *Alb-cre^+/+^* mice. *Fitm2^fl/fl^* mice were used as controls. Mice were maintained on a regular chow diet with *ad libitum* access to food and water at 24°C in a 12:12 h light–dark cycle or, as indicated in Results, at ∼12 weeks of age, mouse had access to a high fat diet (HFD) with 60 kcal% fat (Research Diet, New Brunswick, NJ) for one week prior to analyses of lipoproteins and lipids.

To measure hepatic TG production *in vivo*, we followed our previously published protocol[37], except that instead of TR1339, the non-ionic detergent poloxamer 407 (P-407) was used[38]. Briefly, after baseline plasma samples were obtained, mice were injected intravenously with 1000 mg P-407 per kg body weight (15% vol/vol stock diluted in 0.9% NaCl). One hour after injection, blood samples were collected and plasma was immediately separated. The animals were sacrificed upon completion of the experiment. The livers were flushed with 0.9% NaCl, removed, and homogenized with 1.15% KCl/0.2 mg BHT per ml (9 ml/g tissue). Both the plasma and the liver homogenate samples were quick-frozen in liquid N2 and stored at –80°C until further use. To determine the plasma accumulation of TG after P-407 injection, plasma was collected at the indicated time points and TG was measured by colorimetric assay as described above.

### RNA Isolation and Quantitative Real Time (QRT)-PCR Analysis

RNA was isolated from animal tissues or cultured cells using Trizol Reagent (Life Technologies), following the manufacturer’s instructions. Two μg of total RNA was retro-transcribed to cDNA using the Verso cDNA Synthesis kit (ThermoFisher). RT-PCR was carried using custom-made primers (IDT technologies, Coralville, IA) and the following primers: Rat FITM2 forward primer: CCGAAAGCTACCTCAGCAAC, reverse primer: GTCCGGTGTTCCTGTCTGAT; rat C/EBP homologous protein (CHOP) forward primer: GAAAGCAGAAACCGGTCCAAT; reverse primer: GGATGAGATATAGGTGCCCCC; rat spliced X-box-binding protein-1 (sXBP-1) forward primer: GCTGAGTCCGCAGCAGGT; reverse primer: AGAGGCAACAGCGTCAGAAT; rat Binding immunoglobulin protein (BIP) forward primer: TGGGTACATTTGATCTGACTGGA; reverse primer: CTCAAAGGTGACTTCAATCTGGG; Mouse FITM2 forward primer: GCAACGTCCTCAACGTGTAT; reverse primer: CTGCGATGCTCCTGTCTGAT; mouse CHOP forward primer: CCAACAGAGGTCACACGCAC; reverse primer: TGACTGGAATCTGGAGAGCGA; sXBP-1 mouse forward primer: TGAGAACCAGGAGTTAAGAACACGC; reverse primer: CCTGCACCTGCTGCGGAC; mouse BIP forward primer: TTCAGCCAATTATCAGCAAACTCT; reverse primer: TTTTCTGATGTATCCTCTTCACCAGT. Rat Glyceraldehyde 3-phosphate dehydrogenase (GAPDH) forward primer: TAGTCAACGGGAAAGCCATC, reverse primer: TTCACACCCATGACGAACAT; and mouse 18S rRNA forward primer: GAGGCCCTGTAATTGGAATGAG; reverse primer: CGCTATTGGAGCTGGAATTACC. GAPDH and 16S RNA were used as control genes for the McA cell and liver samples, respectively.

### Microsomal Isolation and ER Purification

Microsomes from liver homogenates were prepared as previously described[39]. Briefly, mice were exsanguinated via cardiac puncture, and livers were removed and washed with TBS. The livers were then homogenized with an electric Heidolph homogenizer (Heidolph Instruments, Schwabach, Germany) at low-medium speed for 10 strokes in homogenization buffer (250 mM sucrose, 20 mM Tris-HCl, pH 7.4, 1 mM EDTA) to yield a 20% (w/v) homogenate. Cell debris was removed by centrifugation at 500xg for 10 min. The supernatant was collected and mitochondria were precipitated by centrifugation at 15,000xg for 10 min, and the supernatant was then centrifuged at 106,000xg for 1 h to obtain a final microsome fraction. The pelleted microsomes were resuspended and washed in 20 nM Tris-HCL, 0.5 M NaCl (pH 7.4) to remove peripherally-associated membrane proteins and LDs, and were then re-pelleted at 106,000xg for 1 h. To release the microsomal content and obtain isolated microsomal (ER) membranes, the purified microsomes were resuspended in alkaline buffer (1 mM Tris-HCl, pH 8.8)[39]. The samples were then incubated on ice for 30 min and centrifuged at 106,000xg for 1 h to pellet the membranes and the supernatant contained the luminal contents. All steps were performed in the presence of protease inhibitors (Sigma-Aldrich).

To isolate ER membranes from McA cells, a modified form of the above homogenization procedure was performed. In brief, cell monolayers were washed with ice-cold TBS and detached, re-suspended, and then homogenized with the same homogenization buffer using a 27G needle-syringe. The purification of ER membranes was then carried out as described above.

### Immunoblot Analysis of Cellular Proteins

Cell and liver lysates were prepared by homogenization in RIPA buffer in the presence of protease inhibitors (Sigma-Aldrich). Total protein amounts were quantified using the Bio-Rad DC assay kit (Bio-Rad, Hercules, CA), and lysate proteins were separated by SDS-PAGE and electroblotted onto PVDF membranes (Bio-Rad). Membranes were blocked with fat-free milk powder (5% w/v) dissolved in Tris-buffered saline containing 0.01% Tween 100 (TBS-T), washed, and incubated overnight at 4°C with rabbit anti-ERp72 (Enzo life sciences, Farmingdale, NY) and anti-GAPDH (Millipore). Secondary antibodies were prepared at a 1:15,000 dilution in TBS-T with 5% fat-free milk powder and incubated for 1 h at room temperature. Goat anti-rabbit conjugated to HRP (BioRad) and donkey anti-goat conjugated to 800CW (Li-Cor) were detected using the Odyssey Fc infrared imaging system (Li-Cor).

### Transmitted Electron Microscopy (TEM)

After density gradient ultracentrifugation of conditioned media (see above), a sample from fraction 1 (which contained VLDL particles) was fixed with 2% paraformaldehyde (Sigma). The particles were visualized using negative staining by placing a small volume onto carbon-coated copper/rhodium grids (Ted Pella Inc., Redding, CA) and staining with 1% uranyl acetate (Polysciences, Inc, Warrington, PA).

ER morphology and lipid droplets were visualized in cultured McA cells fixed with a mixture of glutaraldehyde and paraformaldehyde in 0.1 M sodium cacodylate buffer. To protect intracellular lipid, 1% tannic acid was added to the initial fixation buffer and then incubated in post-fixation solution containing reduced osmium tetroxide for 1.5 h, following established protocols[40]. For the mouse studies, liver tissue samples were washed with water and gradually dehydrated with an ethanol gradient on ice, switched to acetone at room temperature, and embedded in LX112 resin (Electron Microscopy Sciences, Hatfield, PA). For both cells and tissue samples, 70 nm ultra-thin sections were cut and mounted on 200 mesh grids, and 200nm sections were cut and collected on Formvar coated slot copper grids, before staining with uranyl acetate and lead citrate.

All samples were imaged under Talos120C transmission electron microscope (Thermo Fisher Scientific, Hillsboro, OR) using Gatan (4k x 4k) OneView Camera (Gatan, Inc., Pleasanton, CA). Particle diameters were quantified by ImageJ 1.51r software (NIH) based on measurements from 15 images with up to 10 particles per image.

### Simulated Raman Spectroscopy (SRS) Coupled to Fluorescence Microscopy Imaging

McA cells were plated on glass coverslips coated with fibronectin (Sigma-Aldrich) and transfected with plasmids encoding either mCherry-Golgi-7 (Addgene, Watertown, MA), mCherry-ER-3 (Addgene, Watertown, MA), or Perilipin1-YFP[41] following established protocols[32]. The labeled cells were then subjected to SRS to colocalize the presence of intracellular lipids with the Golgi, ER, or LDs markers, respectively. Imaging took place using an inverted microscope (IX81, Olympus, Shinjuku, Japan), as previously described[42].

To image lipids, CH_2_ signals were obtained at 2845 cm^−1^ in the SRS channel. For fluorescence imaging, the same confocal microscope with three continuous visible lasers (405, 488, and 559 nm) controlled by an acoustic optical tunable filter was utilized. Colocalization was established by aligning the fluorescent and SRS channels using fluorescent plastic beads (1 μm diameter, Sigma-Aldrich). The microscope was controlled by Olympus Fluorview 1000 software, and image acquisition was performed by sequentially recording the fluorescence and SRS stacks sequentially. Pearson’s coefficients for fluorescent markers and lipid colocalization measurements were determined using the JACoP ImageJ plugin.

### Confocal Imaging for Neutral Lipid Content

Formalin-fixed livers were sectioned at a 6 μm thickness and mounted onto slides. Samples were stained for 15 min in the dark with BODIPY 493/503 dye (Invitrogen, Carlsbad, CA) diluted in PBS at 1:500 or at 1:1000 for McA cells. Slides/cells were washed with PBS and mounted with ProLong Gold antifade reagent with DAPI (Invitrogen), and confocal images were acquired on a Leica TCS SP5 II confocal microscope (Leica-Camera, Wetzlar, Germany) using a diode laser (excitation 405 nm) and a multiline argon laser (excitation 488 nm) with a 40× Apochromat, numerical aperture 1.25 – Oil objective.

### Lipidomics

Livers were isolated from *Fitm2* knock-out (KO) (*Fitm2^fl/fl^* /*Alb-cre^+/+^*) and “wild-type” (WT; *Fitm2^fl/fl^*) mice at ∼12 weeks of age. Prior to sacrifice, the mice were switched to a high-fat diet (HFD) and were fed for one week. After perfusion with PBS, 100-150 mg pieces of liver from the largest lobes of the livers of 3 mice/group were cut on the day of harvest and immediately flash-frozen in liquid nitrogen and stored at −80°C until further use. At that time, liver samples were powdered and ∼30 mg of tissue was used for extraction. The samples were added to a10-fold volume (based on the weight of the sample) of 80% methanol containing internal standards, and then subjected to bead beating for dispersion. For acyl-coenzyme A quantification, the standards (3-Hydroxyl-3Methylglutaryl-CoA, Acetyl-CoA, Arachidonoyl-CoA, Butyryl-CoA, Glutaryl-CoA, Isovaleryl-CoA, Malonyl-CoA, N-propionyl-CoA, Octanoyl-CoA, Oleoyl-CoA, Palmitoleoyl-CoA, Palmitoyl-CoA, Stearoyl-CoA, Succinyl-CoA) were prepared in the concentration range of 2.3 ng/ml to 556 ng/ml along with the internal standards.

After centrifugation, the supernatant was transferred to a glass vial for LC injection, and the samples were analyzed with a Waters UPLC coupled with ABsciex 6500+ Qtrap-MS system. Metabolite separation was performed on an iHILIC-p column (150mm*2.1mm). Metabolite detection was performed with a Multiple Reaction Monitoring (MRM) mode. A pooled quality control sample was added to the sample list to monitor quality for each metabolite. Mobile phase A contained 10% acetonitrile in water with 20 mM ammonium bicarbonate (pH9.2) and 5 µM PBS, and mobile phase B was 100% acetonitrile. The gradient started with 5% A to 75% A in 15 minutes.

### Statistical analyses

Data are expressed as mean ± standard error of the mean (SEM). Statistical differences were analyzed using GraphPad Prism software (GraphPad Software Inc., San Diego, CA). The distribution normality of sample groups was analyzed using the D’Agostino-Pearson omnibus and the Shapiro-Wilk normality tests. Statistical differences were evaluated by two-tailed Student’s t-testing between groups of two or by one-way ANOVA with Dunnett’s multiple comparison test. Statistical significance was set at P < 0.05. Unless otherwise specified, for cell culture experiments, the results are based on at least 6 wells, and for the mouse studies, the results are based on 3-5 animals.

## Results

### FITM2 deficiency decreases TG content in newly-secreted hepatic lipoprotein particles without affecting particle number

FITM2 promotes cytosolic LD formation in different cell types[43], but its roles in hepatic lipid and lipoprotein metabolism have not been fully characterized. To this end, we began by incubating McA cells, a standard model of mammalian lipoprotein metabolism (e.g., [44]), with *Fitm2* siRNA or control scrambled siRNA. Cells were also incubated with oleic acid (OA)-BSA complexes to stimulate TG synthesis, or with BSA as a control. After dose and time optimizations, we observed that a 72 h incubation with *Fitm2* siRNA reduced *Fitm2* mRNA expression to ∼20% of what was observed with scrambled siRNA (Figure 1A). Using these culture conditions and siRNAs, we then incubated McA cells with [^14^C]-glycerol and [^35^S]-protein labeling mix to quantify newly synthesized TG and apoB100, respectively, in the secreted lipoprotein particles. With both the control and *Fitm2* siRNA treatments, OA increased TG secretion, although this response was significantly attenuated (by ∼45%) with *Fitm2* siRNA (Figure 1B). Because there is only one apoB100 molecule per lipoprotein particle[45, 46], apoB100 recovery in the conditioned medium also estimates the number of secreted particles. Thus, the data in Figure 1C show that in the corresponding treatments in the presence or absence of OA, knocking down *Fitm2* mRNA did not affect apoB100-lipoprotein particle number secretion. In contrast, the data suggest that the secreted particles were lipid-depleted (i.e., denser) when TG synthesis was stimulated and FITM2 was deficient.

**Figure 1.**
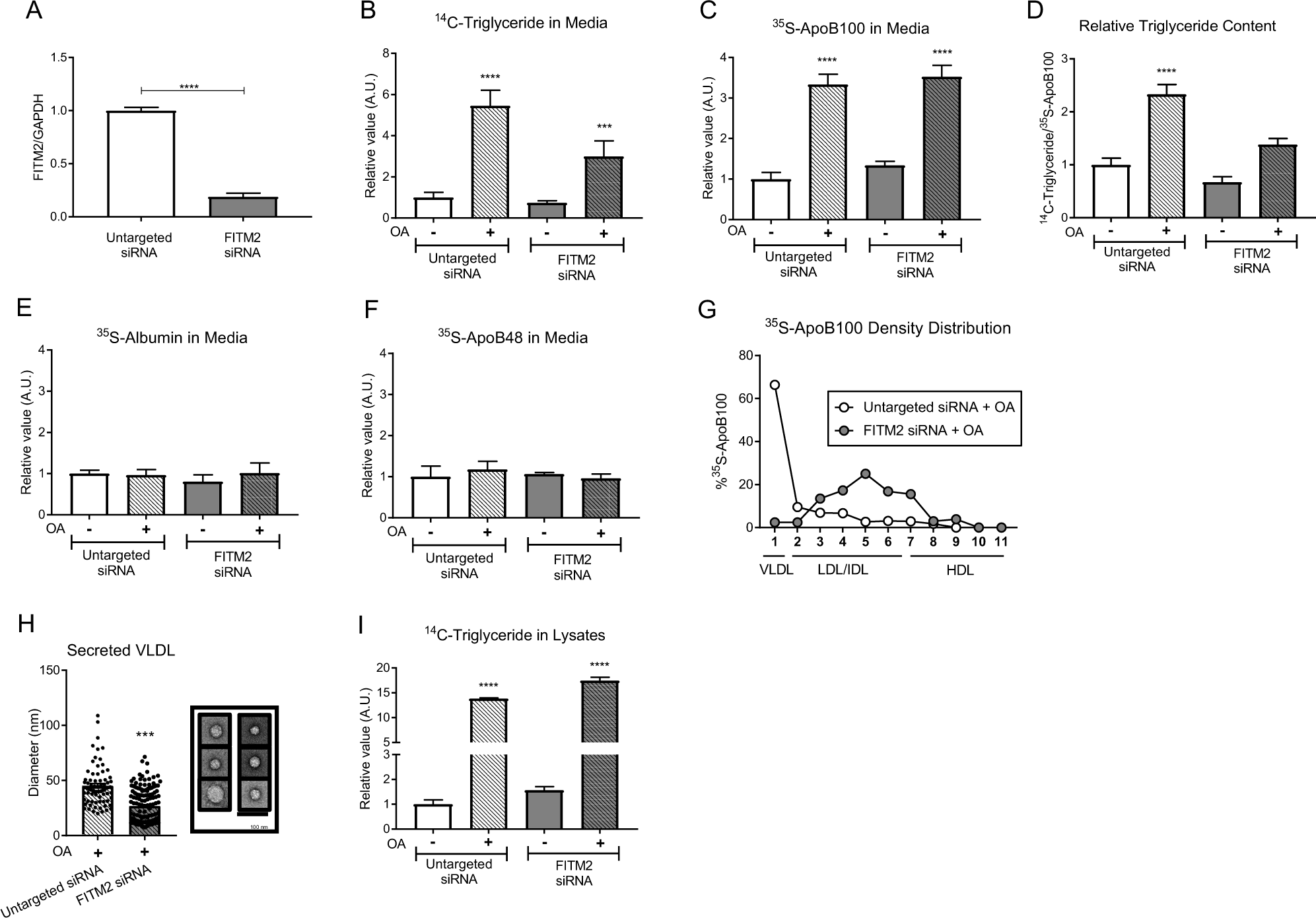
FITM2-deficiency decreases hepatic triglyceride secretion and the density of VLDL particles. **(A)** McA cells were cultured on collagen-coated plates. Cells were treated with siRNA targeting *Fitm2* or untargeted control siRNA for 72 h. *Fitm2* mRNA expression in comparison to *Gapdh* (housekeeping control). **(B)** McA cells were treated with the corresponding siRNA for 72 h and amino acid-starved for 2 h. After this period, cells were incubated with either 0.25 μCi/ml of [^14^C]-glycerol or 120 μCi/ml of [^35^S]-protein labeling mix for 4 h in the presence of OA (+) or BSA alone as a control (−). Conditioned media were collected to determine secreted [^14^C]-TG and **(C)** secreted apoB100 ([^35^S]-apoB100). **(D)** Relative TG content in newly secreted apoB100 lipoprotein particles calculated by dividing [^14^C]-TG by [^35^S]-apoB100. **(E)** [^35^S]-apoB48 and **(F)** [^35^S]-albumin were quantified after phosphorimager analysis. **(G)** Conditioned media samples underwent density-gradient ultracentrifugation to collect 11 individual lipoprotein fractions. [^35^S]-apoB100 quantifications were plotted as the percent of total [^35^S]-apoB100. **(H)** Lipoprotein size was quantified by negative-staining immuno-electron microscopy using the VLDL-like fraction (fraction 1; (**G**)). The particle diameters were measured from 15 images with up to 10 particles per image. Insert shows representative particles from untargeted siRNA (left column) and *Fitm2* siRNA-treated cells (right column). Size quantification was performed with ImageJ. **(I)** Intracellular [^14^C]-triglyceride levels normalized to total [^35^S]-labeled proteins. The experiments were performed in triplicate. All numerical data are presented as mean ± SEM, and statistical differences were evaluated by two-tailed Student’s t-tests or one-way ANOVA. ***, p < 0.001; ****, p < 0.0001.

To test this hypothesis, we measured the ratio between newly-secreted TG and apoB100. As shown in Figure 1D, the ratio was lower in OA-incubated cells treated with *Fitm2* siRNA in comparison to cells treated with control siRNA. Because *Fitm2* siRNA had no significant effect on TG or apoB100 secretion in the absence of OA (Figure 1B,C), these results suggest FITM2 facilitates enhanced lipidation of apoB100-lipoproteins under conditions of increased TG synthesis. Moreover, the specificity of FITM2 deficiency on TG and secreted apoB100 under these conditions was supported by the lack of an effect of *Fitm2* siRNA on the secretion of albumin (Figure 1E) or apoB48 (Figure 1F), a shorter, less lipidated form of apoB that is produced by human and rodent intestinal cells, as well as by rodent hepatic cells[30].

TG synthesized in hepatic cells is primarily secreted via VLDL particles[46]. We hypothesized that a decrease in TG secretion— but no change in apoB100 recovery— meant that the physical characteristics of the lipoproteins in the conditioned medium were altered. Thus, we characterized apoB100-containing particles in conditioned media from cells treated with OA using two independent approaches. First, the majority of newly-secreted apoB100 particles were detected in buoyant density gradient fractions that contain lipoproteins of VLDL density (i.e., fraction 1, Figure 1G) when assayed from control siRNA-treated cells, as expected[47]. In contrast, after *Fitm2* siRNA treatment the majority of the apoB100 particles were found in denser fractions (i.e., fractions 3-7, Figure 1G). These data indicate that FITM2-depleted cells exhibit a reduced capacity to fully lipidate nascent VLDLs. The data are also consistent with the concept that despite the decrease in TG associated with secreted apoB100-lipoproteins when FITM2 was knocked down, there was still sufficient early lipidation of apoB100 by MTTP. This would spare the nascent particles from ER-associated proteasome-mediated degradation[48], a hypothesis that is also consistent with the fact that the number of secreted particles was unaltered by FITM2 deficiency (Figure 1C).

In the second approach, we quantified the diameters of the lipoproteins in the VLDL-containing fraction (fraction 1) using transmission electron microscopy (TEM; Methods). *Fitm2* siRNA-treated McA cells secreted significantly smaller lipoproteins in the VLDL fraction in comparison to those secreted from control siRNA-treated cells (Figure 1H), consistent with less TG per VLDL particle. This is the fourth independent indicator— along with altered TG secretion, the TG/apoB100 ratio, and the gradient densities— that FITM2 is essential to fully lipidate apoB100-lipoproteins when TG synthesis is stimulated.

To further support this conclusion, we confirmed that TG synthesis was not decreased when FITM2 was knocked down, an outcome that might contribute to reduced lipid loading. Indeed, decreased TG synthesis was previously observed in an immortalized human breast carcinoma cell line when FITM2 was deficient[28]. Therefore, we measured the accumulation of newly synthesized TG in control and *Fitm2* siRNA-treated cells in the presence or absence of OA. As shown in Figure 1I, OA stimulated accumulation of newly synthesized TG in both control and *Fitm2* siRNA-treated McA cells, as anticipated, but interestingly FITM2 depletion modestly increased TG levels. Thus, *Fitm2* siRNA-treatment allows for ongoing TG synthesis, but TG-loading onto VLDL particles is compromised.

### TG accumulates in the ER in FITM2-deficient cultured hepatocytes

FITM2 deficiency resulted in secreted apoB100-lipoprotein particles that were smaller and denser (i.e., lipid-depleted), in spite of a modest increase in total cellular TG content (Figure 1). To determine the site(s) of TG accumulation, we first used confocal fluorescent microscopy to visualize three relevant lipid-rich cellular locations. This analysis was then combined with simulated Raman spectroscopy (SRS) microscopy to examine the location of neutral lipids, which are mostly TG after treatment with OA[49]. As shown in Figure 2A, the Golgi apparatus (“mCherry Golgi-7”), the ER (“mCherry-ER-3”), and LDs (“PLIN1-YFP”) were readily detected, as was the location of neutral lipids. In addition to being examined in the presence or absence of OA, we also examined the McA cells under conditions in which FIT2M was unaffected or depleted (using control or *Fitm2* siRNA, respectively). After image quantification, Pearson’s coefficients were calculated. The low Pearson’s coefficients indicated that neutral lipids did not strongly co-localize with the Golgi (i.e., mCherry-Golgi-7), and any residence in this compartment was independent of the siRNA treatment or OA (Figure 2B). In contrast, McA cells treated with control siRNA showed less colocalization between neutral lipids and the ER marker, mCherry-ER-3, than those cells treated with *Fitm2* siRNA (Figure 2C), and this was independent of OA addition. Instead, there was greater colocalization with the LD marker, PLIN1-YFP[10], in control-treated cells versus those treated with *Fitm2* siRNA. This was again independent of stimulated TG synthesis (Figure 2D). Together, these data suggest that hepatic FITM2 deficiency shifts the relative distribution of TG from LDs to the ER, and that FITM2 plays a role in TG distribution under both homeostatic and lipid-stimulated conditions.

**Figure 2.**
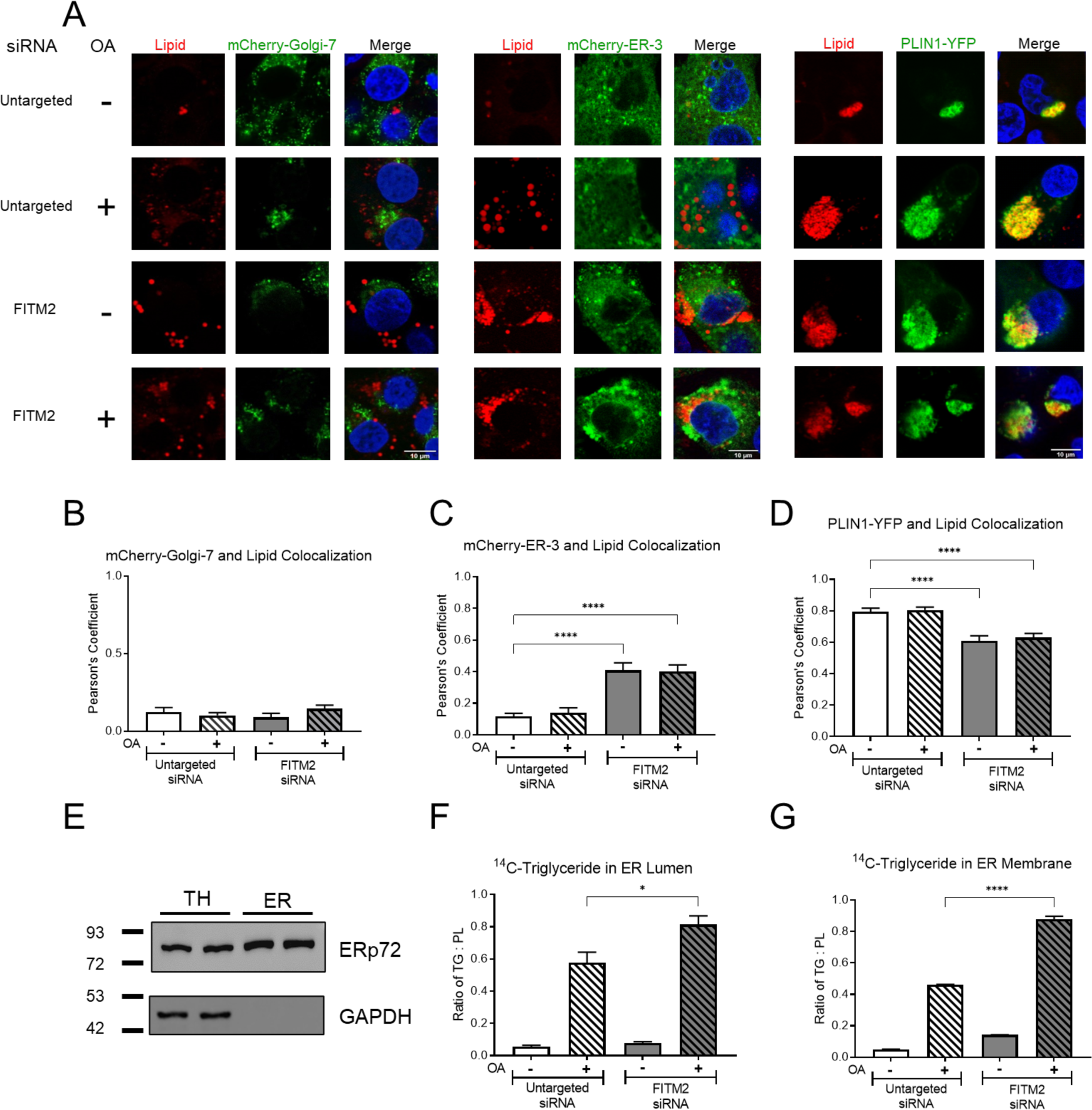
FITM2-deficiency promotes lipid accumulation in the ER of cultured hepatic cells. **(A)** McA cells were cultured on fibronectin-coated glass coverslips. Cells were treated for 72 h with siRNA targeting *Fitm2* or untargeted control siRNA. After 48 h, McA cells were transfected with either mCherry-Golgi-7, mCherry-ER-3, or perilipin 1 (PLIN1)-YFP expression plasmids to label the Golgi apparatus, the ER, or cytosolic lipid droplets, respectively. After 24 h, cells were washed with PBS and incubated for 4 h with media supplemented with 0.4 mM of oleic acid (OA) conjugated with BSA (+) or unconjugated BSA as control (−). The cells were fixed with 10% formalin and mounted for visualization. Simulated Raman Spectroscopy (SRS) was utilized to visualize intracellular lipids, which are mostly triglycerides (red), while confocal microscopy was utilized to visualize fluorescently-labeled proteins (green) and DAPI-stained nuclei (blue). **(B)** Pearson’s colocalization indices between lipids and mCherry-Golgi-7, **(C)** mCherry-ER-3, and **(D)** PLIN1-YFP. The extent of colocalization was measured by analysis of regions in yellow using the ImageJ Pearson’s coefficient tool. **(E)** McA cells were subjected to siRNA targeting *Fitm2* or untargeted (control) for 72 h. Cells were then incubated with [^14^C]-glycerol and [^35^S]-protein mix for 4 h in the presence of OA (+) or BSA alone (−). Cells were harvested and ER membranes isolated (Methods). The enrichment of an ER marker shown was measured by western blot in comparison to the total cell homogenate (TH) using ERp72 and GAPDH, respectively. After isolation of the respective compartments, **(F)** the [^14^C]-TG content was determined in the ER lumenal fraction, **(G)** and the ER membrane fraction. Samples were normalized to total [^35^S]-labeled proteins in the cell lysate. Experiments were performed in triplicate. Colocalization experiments were performed using at least 10 cells for each experiment. Numerical data are presented as mean ± SEM, and statistical differences were evaluated by one-way ANOVA. ***, p < 0.001 ; ****, p < 0.0001 versus untargeted siRNA, (+OA, -OA).

To independently confirm these data and examine absolute changes in the distribution of newly-synthesized TG, we next quantified TG content in the ER membrane by incubating control and *Fitm2* siRNA-treated McA cells with [^14^C]-glycerol in the presence or absence of OA, as in Figure 1. We then purified microsomal membranes, which were highly enriched in ER as shown by the recovery of the protein ERp72 (Figure 2E). We then determined the [^14^C]-TG content in this fraction by thin layer chromatography. Microsomes were also used to prepare the lumenal contents, and [^14^C]-TG was measured in the membrane and lumenal fractions. This analysis showed that OA-stimulated TG synthesis increased [^14^C]-TG content in both the ER lumen (Figure 2F) and membrane (Figure 2G). In addition, treatment with *Fitm2* siRNA led to a greater accumulation of TG in these locations. Overall, these results indicate that FITM2-deficency reduces cytosolic LD, but ER lumenal and membrane-resident TG increase when cellular TG accumulates. More generally, these findings indicate that the incomplete lipidation of VLDL particles in FITM2-deficent cells (Figure 1) was not a result of a decrease in TG availability, but more likely reflects defective coupling between TG and VLDL assembly in the ER.

### ER-resident TG also accumulates in liver-specific FITM2-deficient mice

To examine whether FITM2 deficiency also alters hepatic TG and apoB100-lipoprotein metabolism *in vivo*, we generated a liver-specific FITM2-deficient mouse by crossing *Fitm2*-floxed mice (*Fitm2^fl/fl^*) with mice overexpressing cre-recombinase under the regulation of the albumin promoter (*Alb-cre^+/+^*). *Fitm2^fl/fl^*/*Alb-cre^+/+^* mice lacked any obvious signs of developmental abnormalities (data not shown), despite expressing marginal levels of hepatic *Fitm2* mRNA in comparison to *Fitm2^fl/fl^* control mice (Figure 3A).

**Figure 3.**
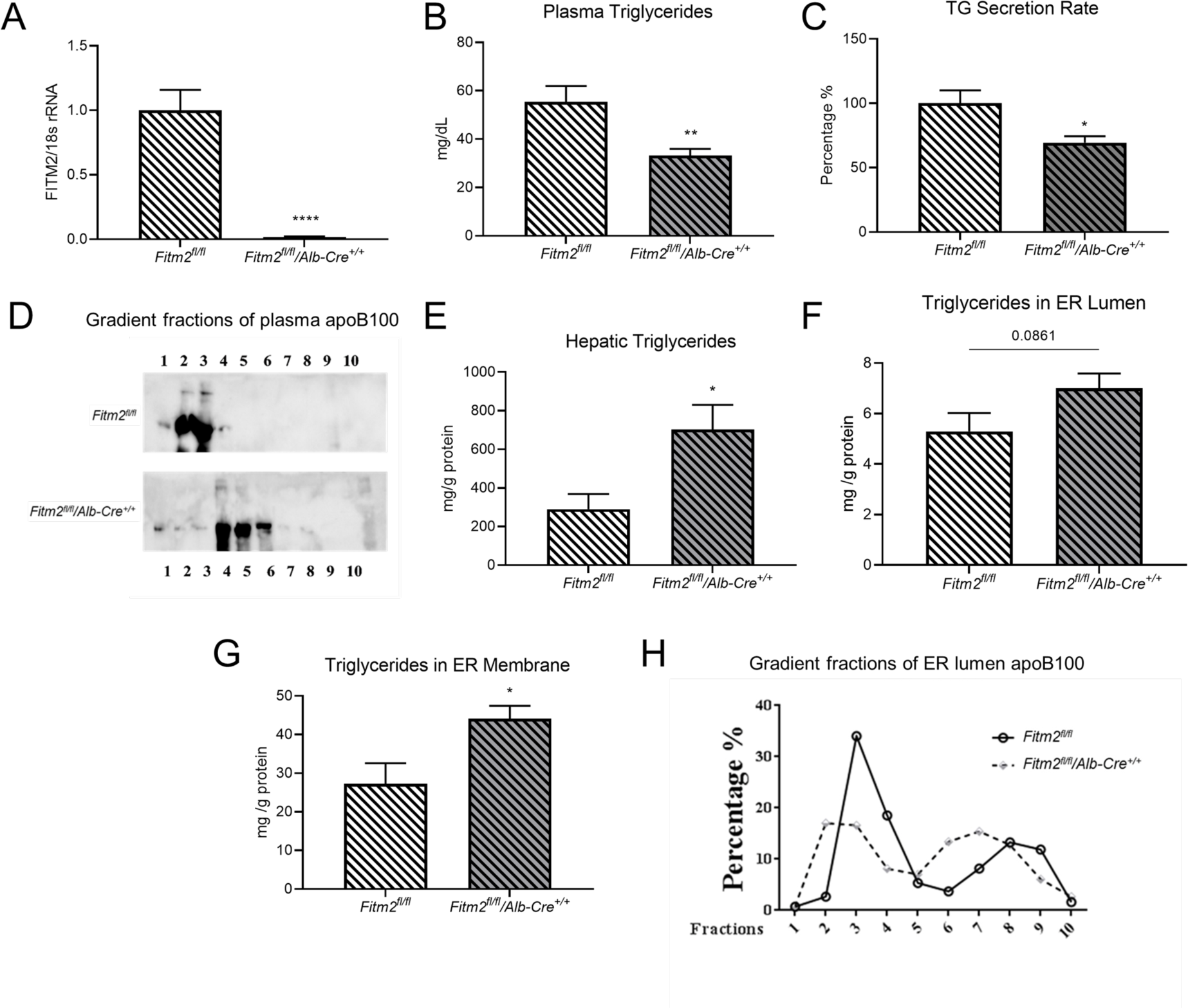
Effects of liver-specific FITM2-deficiency on plasma and hepatic triglyceride metabolism and apoB100-lipoprotein density. Four-week old age and sex matched and *Fitm2^fl/fl^/Alb-cre^+/+^* mice were fed HFD for one week. **(A)** *Fitm2* mRNA expression levels in liver homogenates, and **(B)** Plasma TG levels are shown. **(C)**Hepatic TG secretion rates, represented as a percentage of those in the *Fitm2^fl/fl^* mice, were measured after poloxamer 407 injection as described in the Methods. **(D)** A representative western blot of apoB100 across density gradient fractions of plasma lipoproteins from the top/lowest density (fraction 1) to the bottom/highest density (fraction 10). **(E)** Total hepatic TG levels were measured and ER microsomes were isolated and separated to resolve the levels in the **(F)** lumenal and **(G)** membrane fractions. **(H)** The isolated ER lumenal contents were resolved by density gradient ultracentrifugation, as in **(D)**, and the apoB100 content of each fraction was determined by western blotting. Numerical data are presented as mean ± SEM of 3 to 8 animals/group, and statistical differences were evaluated by one-way ANOVA or by two-tailed Student’s t-test. *, p < 0.05; **, p < 0.01; ****, p < 0.0001 versus *Fitm2^fl/fl^*mice fed HFD.

Since the loss of FITM2 *in vitro* affects the characteristics of secreted VLDL (Figure 1), we first examined plasma TG levels in control and FITM2-deficient mice. To this end, we fed sex and age-matched *Fitm2^fl/fl^*/*Alb-cre^+/+^* and *Fitm2^fl/fl^* (control) mice a high-fat diet (HFD) for one week to stimulate hepatic TG synthesis, then harvested tissues and plasma in the fed state (analogous to acutely supplying OA *in vitro*). Under these conditions, plasma TG levels were lower in *Fitm2^fl/fl^*/*Alb-cre^+/+^*mice (Figure 3B), with the changes by FPLC mainly in the VLDL fraction (data not shown). These results are consistent with the *in vitro* data that FITM2 function is particularly vital when abundant levels of TG are available to lipidate nascent VLDL particles.

This suggestion was investigated further by an *in vivo* hepatic TG production study in which the rate of secretion can be directly measured (Figure 3C). In the HFD fed state, TG production was lower in *Fitm2^fl/fl^*/*Alb-cre^+/+^ (i.e., liver*-deficient) mice, compared to control mice. Also in agreement with the McA cell results, apoB100-containing lipoproteins in the plasma of the *Fitm2^fl/fl^* /*Alb-cre^+/+^* mice were more dense (i.e., lipid-depleted), as shown by their shift to fractions of higher density after ultracentrifugation (Figure 3D). This was concomitant with increased hepatic TG content (Figure 3E), as well as an increase in TG content in the lumen or membrane of hepatic ER-derived vesicles (Figure 3F and Figure 3G, respectively).

To better define the nature of the apoB100-containing lipoproteins that resided in the ER lumen, the particles were fractionated using density centrifugation. While the percentage of lipoproteins in the denser fractions (fractions 6-10) was similar, the net amounts of apoB100-containing lipoproteins in the more lipidated fractions (fractions1-5) decreased. This result is generally consistent with the cell-culture results (Figure 1). One model to explain these data is that, as noted previously, the early lipidation of apoB100, which is mediated by MTTP[50], occurs normally, which allows apoB100 to escape proteasome-dependent degradation[51]. Based on the data *in vitro* and *in vivo*, the subsequent addition of TG to the nascent lipoprotein (referred to as “second step” lipidation[52]) to form more buoyant VLDL particles is impaired when FITM2 is deficient.

### FITM2 deficiency promotes hepatic ER stress in vitro and in vivo

Accumulation of lipids in the ER is known to trigger ER stress and the expression of genes associated with the unfolded protein response (UPR), which if unmitigated compromises homeostasis and can trigger cell death[53, 54]. Given that FITM2-deficency increases the content of ER TG *in vitro* and *in vivo* (Figures 2 and 3), we hypothesized that this would induce the UPR. Thus, we next examined the levels of mRNAs associated with this pathway[55, 56]. In McA cells treated with control or *Fitm2* siRNA, the levels of several UPR targets increased in the presence of OA: the ER Hsp70 molecular chaperone, BiP, spliced XBP-1 (sXBP-1), which is the most proximal reporter for activation of one arm of the UPR[57, 58]), and CHOP (a downstream pro-apoptotic factor) were elevated in control cells treated with OA; these levels rose even higher when FITM2 was depleted (Figure 4A-C, respectively). We then asked if UPR induction was also triggered in mice lacking hepatic FITM2. For this experiment, control and FITM2-deficient mice were maintained on standard chow or were switched to a HFD diet for one week before the levels of each UPR reporter was measured in liver samples. Interestingly, effects on the UPR were largely unchanged in control mice in the presence or absence of a HFD, perhaps because UPR-induced regulation of lipid synthesis supports the return to an unstressed homeostatic state[59]. The combination of FITM2-deficiency and stimulation of TG synthesis as a result of the HFD, however, significantly elevated the stress-related mRNAs (Figure 4 D-F).

**Figure 4.**
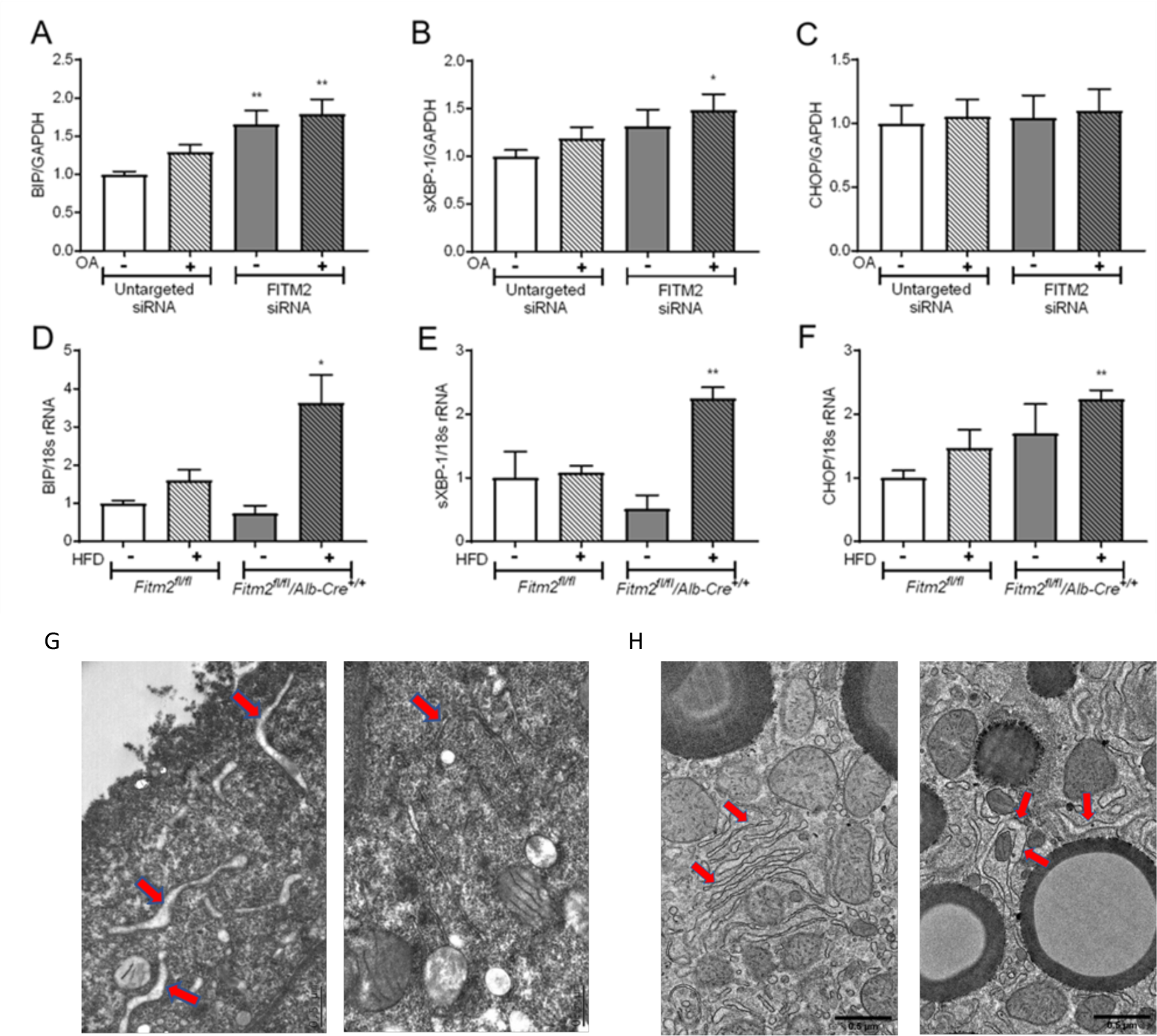
Hepatic FITM2 deficiency in lipid-loaded hepatic cells results in ER stress both *in vitro* and *in vivo*. **(A-C)** McA cells were cultured on collagen-coated plates, and cells were treated with siRNA targeting *Fitm2* or an untargeted control siRNA for 72 h. The McA cells were exposed to 0.4 mM of oleic acid (OA) conjugated with BSA (+) or unconjugated BSA as a control (−) for 4 h before cells were harvested to examine mRNA expression of ER stress-associated markers by RT-PCR. **(D-F)** Four-week old, age and sex matched *Fitm2^fl/fl^* and *Fitm2^fl/fl^*/*Alb-cre^+/+^*mice were fed a standard diet (−) or HFD (+) for one week. Mice were fasted overnight before harvesting the liver to measure hepatic mRNA levels of the ER stress markers. **(G)** EM images show numerous examples of swollen ER structures (red arrows) in FITM2-deficient McA cells incubated with OA (left panel), but not in FITM2-sufficient cells similarly treated (right panel). (**H**) Four-week old age and sex matched *Fitm2^fl/fl^* and *Fitm2^fl/fl^/Alb-cre^+/+^* mice were fed a HFD for one week. The mice were perfused with 4% paraformaldehyde in PBS, and liver samples were prepared to visualize ER structures. The red arrows highlight the ER. Statistical differences in panels A-F were evaluated by one-way ANOVA *, p < 0.05; **, p < 0.01.

Because ER stress can also lead to ER expansion and altered ER morphology, we next performed transmission electron microscopy (TEM) in McA cells and mouse livers. McA cells treated with *Fitm2* or control siRNA and incubated with OA to promote TG synthesis displayed altered ER structure, consistent with studies in the breast carcinoma cell line model[28]). Expansion was evident by a dilated ER when *Fitm2* siRNA-treated McA cells were incubated in the presence of OA (Figure 4G). Similar morphological ER alterations in liver-specific FITM2-deficient hepatocytes from mice fed a HFD for one week were also evident (Figure 4H).

Another possible contributor to ER stress was proposed by recent reports by Farese and Walther[5, 28]: the acyl-CoA diphosphatase activity associated with FITM2 were proposed to be required to maintain ER membrane homeostasis. More specifically, they suggested that FITM2 deficiency remodels the phospholipid composition of the ER membrane, which subsequently results in ER stress. They also reported that a number of acyl-CoA substrates accumulated in FITM2-deficient mice fed a chow diet[5]. Consequently, we performed a lipidomics analysis of long chain-acyl-CoA derivatives from control and *Fitm2^fl/fl^*/*Alb-cre^+/+^* mouse livers to determine if a similar phenomenon occurred. Given the robust effects of hepatic FITM2 deficiency on VLDL lipid assembly and secretion with the HFD (Figure 3), we used liver samples from these mice.

The data shown in Table 1 are consistent with the results of Bond et al.[5] in that the levels of four prominent long-chain fatty acyl-CoA derivatives in the liver rose significantly when FITM2 was deficient. Specifically, the measured levels of palmitoyl-CoA, palmitoleoyl-CoA, steroyl-CoA, and oleoyl-CoA all significantly increased between 2.0- and 3.6-fold when the *Fitm2^fl/fl^* /*Alb-cre^+/+^* mice were compared to control littermates. In the future, it will be important to determine the extent to which the ER stress-associated phenotypes we observed in the FITM2-deficient animals (Figure 4) derive from altered levels of specific CoA derivatives versus the noted accumulation of lipids in the ER membrane (Table 1 and[5]). Regardless, it is clear that the loss of hepatic FITM2 exhibits profound effects on lipid homeostasis *in vitro* and *in vitro*, and—as reported here for the first time—on the accumulation in hepatocytes of ER lipids and TG loading onto nascent apoB100-containing VLDLs.

**Table 1.** The loss of hepatic FITM2 increases levels of long chain fatty acyl-CoA derivatives. Mice were switched to a high fat diet (HFD) for one week and liver extracts were prepared for lipidomic analysis, as described in Methods. Data for each mouse/genotype are shown as individual values as well as averages of the three mice in each group. Units are pg/ml. P-values were calculated using a two-tailed Student’s -test.

| Fatty Acyl-CoA identity | Fatty Acid component | FITM2 KO values (units) | FITM2 KO mean (units) | WT average (units) | WT mean (units) | Fold Change (KO/WT) | Statistics (P-value) |
| --- | --- | --- | --- | --- | --- | --- | --- |
| Palmitoyl-CoA | 16:0 | 0.085, 0.086, 0.101 | 0.091 | 0.040, 0.066, 0.010 | 0.039 | 2.3 | 0.038 |
| Palmitoleoyl-CoA | 16:1 | 0.086, 0.092, 0.106 | 0.095 | 0.018, 0.055, 0.006 | 0.026 | 3.6 | 0.013 |
| Steroyl-CoA | 18:0 | 0.137, 0.147, 0.157 | 0.147 | 0.048, 0.122, 0.028 | 0.066 | 2.2 | 0.049 |
| Oleoyl-CoA | 18:1 | 0.247, 0.244, 0.272 | 0.254 | 0.068, 0.154, 0.029 | 0.083 | 3.0 | 0.010 |

## Discussion

The assembly of a VLDL particle entails a complex choreography of events that are only partially understood. These events include the independent syntheses in the smooth ER of multiple lipids (e.g., sterols, triglycerides, and most phospholipids) as well as the synthesis in the rough ER of its essential protein component, apoB100. The initial formation of a nascent VLDL particle (sometimes referred to as pre-VLDL) requires the transfer by microsomal triglyceride transfer protein (MTTP) of the lipid components to the translocating apoB100 polypeptide. In situations in which there is insufficient lipidation, due to either inadequate lipid availability or MTTP activity, apoB100 is unstable and recognized for ER-associated degradation (ERAD), which shunts immature apoB100 to ER-associated ubiquitin ligases and then the cytosolic proteasome for degradation (see [48] for a review). If instead, MTTP-dependent lipid-loading is adequate, further lipidation of the nascent particles in the ER occurs, and several factors have been implicated in this process, including PLIN2[15], CIDEB[15], LPCAT3[16], apoC3[17], seipins[18], and Tm6sf2[19–21]. Based on the data presented in this report and by others[5], FITM2 now claims a place on this list.

FITM2 was initially discovered as an integral smooth ER membrane protein required for cytosolic LD formation[10], with its function generally conserved from yeast to mammals (reviewed in[60]). In addition to facilitating cytosolic LD formation, FITM2 was also proposed to participate in VLDL lipidation in liver[29]: FITM2 was speculated to mediate TG transfer bi-directionally; i.e., to both the cytosol and the ER lumen to respectively play roles in nascent cytosolic LD or ER lumenal VLDL particle biogenesis.

In support of this speculation, we now report that: 1) FITM2-deficient hepatic cells *in vitro* or *in vivo* secrete TG-depleted VLDL particles, but the number of particles is unchanged compared to controls; 2) FITM2 deficiency in mice on a HFD decreases plasma TG levels, with the number of apoB100-containing lipoproteins remaining similar, but shifting from VLDL to LDL density; 3) When TG synthesis is stimulated and FITM2 is deficient, TG accumulates in the ER both *in vitro* and *in vivo*. Despite its availability, this TG pool is unable to fully lipidate apoB100 particles, and, 4) FITM2 deficiency disrupts ER morphology and results in ER stress. Each of these results is discussed in more detail below.

That FITM2 deficiency reduces VLDL-TG content in hepatic cells could not be explained by decreased TG synthesis (Figure 1), which was observed in FITM2-deficient breast carcinoma cells[28]) or by decreased MTTP activity (data not shown). Rather, our data support FITM2’s role as an apoB100/VLDL lipidation factor likely acting after MTTP, given the insignificant changes in apoB100 secretion when FITM2 is depleted *in vitro* (Figure 1) and *in vivo* (Figure 3), and evidence (reviewed in[48]) that if the MTTP step is not successful, nascent apoB100 polypeptides are targeted for ERAD. Further supporting FITM2 as a lipidation factor is our finding that within the ER lumen, the shift of apoB100-containing lipoproteins from dense to lighter (i.e., more lipidated) fractions is impaired when TG synthesis is stimulated (Figure 3H).

Since TG synthesis increases—yet TG secretion decreases—when FITM2 is depleted in hepatic cells and liver, we suspected that TG accumulates intracellularly. This was conclusively established based on multiple lines of evidence. First, TG measurements were made on cell or tissue lysates (Figures 1 and 3, respectively), and by using two independent methods (SRS microscopy and subcellular fractionation) we localized the accumulated lipids to the ER, consistent with studies in yeast[9], *C. elegans*[61], zebra fish[10], and mouse intestine[62]. Second, we noted structural evidence of TG accumulation in the ER in EM images obtained from FITM2-deficient hepatic cell lines and liver samples (Figure 4). Though ER morphology was unchanged with FITM2-deficiency in chow-fed mice[5], alterations in ER structure have been described in breast carcinoma cells[28], yeast[9], and mouse enterocytes[62] when FITM2 or the FITM2 homolog(s) were absent. In earlier studies in yeast and mouse enterocytes, lipid accumulation was associated with morphological changes, as we observed (Figure 4). In the *in vitro* and *in vivo* analyses reported here, TG also accumulated in the ER membrane along with the lumen in the FITM2-deficient state (Figures 2 and 3). FITM2-deficiency similarly resulted in ER membrane lipid accumulation in yeast[9] (and reviewed in[60]), whereas LDs were found in the ER lumen in mouse intestinal cells[62].

Because ER lipid accumulation induces ER stress (e.g.,[54], we investigated whether the same phenomenon arises in FITM2-deficient hepatic cell lines and mouse liver. As shown in Figure 4, several well-established UPR markers were increased, consistent with results from other studies (e.g., [5, 28, 63]). While it is likely that this results from an inability to maximally export ER TG via VLDL particles, which is compounded by an inability to fully transfer ER-derived TG to cytosolic LDs—an independent inducer of ER-stress[64]—it has also been proposed that FITM2’s enzymatic activity maintains ER membrane homeostasis[5, 28]. While our future efforts will seek to address the importance of FITM2 enzymatic activity on cellular and organismal lipid homeostasis, especially when animals are fed a HFD (see below), we note that the levels of four long chain fatty acyl-CoA derivatives were significantly higher in the livers of FITM2-deficient animals (Table 1), which is consistent with defects in the enzymatic degradation of these lipids.

While this report was in preparation, another report on the role of FITM2 in mouse liver was published (Bond et al.,[5]). In contrast to the present studies, most effects of hepatic FITM2 deficiency were conducted in mice on a chow (i.e., low fat) diet. We instead examined mice on a HFD, which is more clinically relevant given that HFDs are typically consumed in the setting of NAFLD that is found in ∼70% of overweight people[65]. Indeed, there was a greater impact of FITM2 deficiency on TG secretion when lipid synthesis in hepatic cells was stimulated (compare +/− OA columns in Figure 1B). Consistent with these *in vitro* results, hepatic FITM2 deficiency had significantly more pronounced effects on TG production *in vivo* in the fed state (compare Figure 3C vs. Supplemental Figure 1), as well as the UPR (Figure 4D-F). Together, our *in vitro* and *in vivo* data indicate that FITM2 plays a more pronounced role on ER lipid homeostasis and VLDL lipidation when there is a surfeit of newly-synthesized hepatic TG, a situation frequently manifested in cardiometabolic diseases (e.g., obesity and insulin resistance).

In chow-fed mice, Bond et al. reported increased TG content in the livers of FITM2-deficient mice, along with an increase in the levels of ER stress markers[5]. They also reported smaller VLDL particles in circulation, but plasma levels of TG were unaltered. In both our work and in the limited studies in Bond et al.[5], plasma TG levels were lower in mice on a HFD. We found, however, that hepatic TG content increased, whereas Bond et al. saw decreased hepatic TG content. One explanation for this discrepancy is the length of feeding on the HFD. We chose to perform all measurements after one week of HFD feeding. Under these conditions, the weights of both the FITM2-deficient and sufficient mice rose normally by ∼2 grams, and the animals exhibited simple steatosis (data not shown), similar in histological appearance to NAFLD in humans (e.g., [66]). In contrast, Bond et al.[5] fed mice a HFD for 11 weeks, so the animals exhibited a number of adverse effects, such as poor weight gain and multiple indices of liver injury (e.g., histological changes and increased plasma levels of transaminases). We suggest our results are not confounded by the apparent toxicity of the combination of chronic FITM2-deficiency and consumption of a long-term HFD, thereby better reflecting the physiological role(s) of FITM2.

In summary, the present results suggest a model in which FITM2 contributes to VLDL lipidation, especially when newly synthesized hepatic TG is in abundance. As noted above, FITM2 likely acts in the ER after the initial lipidation by MTTP of translocating apoB100 polypeptides. It was previously shown that post-MTTP lipidation of pre-VLDL within the ER occurs while apoB100 remains associated with the inner leaflet of the smooth ER membrane (reviewed in[52]), the site at which FITM2 and DGAT2 (which catalyzes the final enzymatic step to form TG) reside. Increases in ER membrane TG and lumenal LDs reflect both a role for FITM2 in mobilizing newly synthesized lipids to the ER lumen, as well as facilitating the transfer of TG from lumenal LD to pre-VLDL (see[67] for a review of this process). Ultimately, further studies will be needed to test the specific contribution of FITM2 on “second-step” ER lipidation and to determine 1) the relationship between FITM2 and the actions of the other factors that promote pre-VLDL lipidation (see above); 2) the role of FITM2 in NAFLD and NASH, and, 3) the basis for ER stress/UPR in the FITM2-deficient state. Nonetheless, our studies provide a new conceptual framework that will guide future investigations.

## Acknowledgements

Funding sources for the authors (identified by their initials) are the following: HW, CN, MLF, EAF: NIH R01 HL168569, P01 HL160470; JLB: NIH R01 HL168569, P01 HL160470, NIH GM131732; JA: NIH R01 HL168569, R01 HL147252, P01 HL160470; YY, MCW: Howard Hughes Medical Institute; YQ, MC, IJK: NIH P60DK020541; IJG: NIH P01 HL160470; MMH: NIH R01 HL158054, R01 HL166214, P01 HL160470; SR: American Heart Association 19POST34410063, NIH P01 HL160470. IJK notes that the mass spectrometry assessments were done on a Sciex 6500+ QTRAP funded by a S10 SIG Award (1S10OD021798).

We thank NYU Langone Health DART Microscopy Lab Alice Liang and Kristen Dancel-Manning for their assistance with TEM work. This core is partially funded by NYU Cancer Center Support Grant NIH/NCI P30CA016087.

---

**Supplemental Figure 1.**
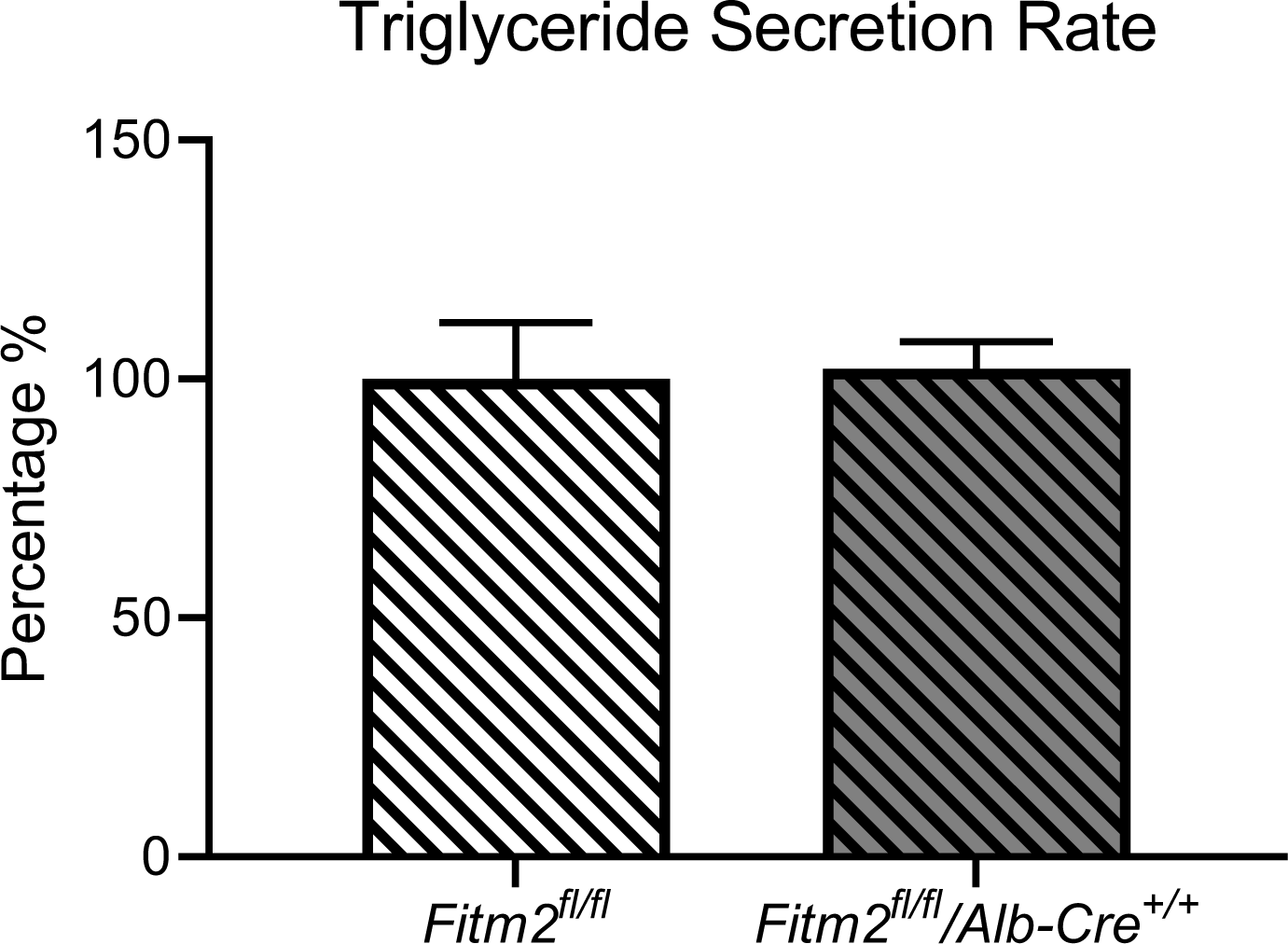
The secretion of triglyceride is unchanged in fasted *Fitm2^fl/fl^* and *Fitm2^fl/fl^*/*Alb-cre^+/+^* mice. The TG secretion rate as a percentage of the control in the indicated mice was performed as in Figure 3C, except that the mice were in the fasting state. There was no significant difference as assessed by a two-tailed Student’s t-test.

